# Repeated Binge-Like Alcohol Drinking Heightens Aggression in Mice

**DOI:** 10.1101/2025.04.22.650080

**Authors:** Micah D. Frier, Nicole P. Biggi, Jessica A. Babb, Emily L. Newman, Herbert E. Covington, Marcus M. Weera

**Affiliations:** Department of Psychology, Tufts University, Medford, MA; Department of Neuroscience, School of Medicine, Tufts University, Boston, MA, USA; Department of Psychiatry, Harvard Medical School McLean Hospital, Belmont, MA, USA; Department of Psychology and Human Development, SUNY Empire State University, Saratoga Springs, NY, USA; Neuroscience Program, Graduate School of Biomedical Sciences, Tufts University, Boston MA

**Keywords:** alcohol, aggression, abstinence, Drinking in the Dark, neutral arena, resident-intruder test, CFW mice, binge drinking

## Abstract

**Rationale:** In humans, alcohol drinking is a significant driver of violent behaviors such as assaults and homicides. While acute intoxication is known to produce heightened aggression, little is known about alcohol’s long-term effects. Emerging evidence, however, suggests that chronic alcohol intake can promote heightened aggression, including during abstinence and may also sensitize individuals to alcohol’s acute aggression-heightening effects.

**Objectives:** The goal of this study was to test the effects of chronic binge-like ethanol drinking on both alcohol-involved and alcohol-uninvolved aggression in male CFW mice. We aimed to model individual differences in binge drinking and assess changes in aggression during both acute and protracted abstinence.

**Results:** After 5 weeks of Drinking in the Dark (DID), CFW mice that showed higher levels of EtOH drinking (‘high drinkers’, 1.33 g/kg/h) became more aggressive than low drinkers (0.45 g/kg/h) and H₂O controls, as measured via frequency of attack bites during resident-intruder fighting. In the first aggressive encounter following 1 week of abstinence, animals with an alcohol drinking history initiate a fight more rapidly and with greater consistency than H₂O controls. We also found that a single session of binge-like alcohol drinking acutely heightened aggression regardless of drinking history.

**Conclusions:** These results suggest that repeated binge-like alcohol drinking causes escalations in alcohol-uninvolved aggression during acute (in high drinkers) and protracted abstinence (in all alcohol drinkers). However, chronic alcohol intake does not appear to sensitize animals to alcohol-involved aggression. These findings support the utility of genetically heterogeneous CFW mice for modeling individual variability in alcohol-related aggression.

## Introduction

Approximately half of all violent crimes committed in the United States are associated with alcohol intake (Beck et al., 2014). It has been posited that alcohol has both acute effects, causing heightened aggression immediately after drinking, as well as chronic effects, wherein chronic alcohol intake causes long-term changes in aggressive behavior (Lipsey et al., 1997; Miczek et al., 2015; Robertson et al., 2020; White et al., 2009). Acute effects have been thoroughly studied and well characterized, whereas chronic effects are complex and debated. It is well known that acute alcohol intoxication can cause heightened aggression in humans (Duke et al., 2015) and in rodents (Miczek et al., 1992, 1998). Reliable and reproducible methods have been developed to study acute alcohol-heightened aggression in mice, rats and monkeys, and the effects of alcohol intoxication have been repeatedly demonstrated across species (Miczek et al., 2002, 2004). In mice, acute intoxication causes not only an increase in bite frequency in resident-intruder paradigms, but also more extreme forms of aggressive behavior (Newman et al., 2018), as well as an escalation in the motivation to engage in a fight in operant paradigms (Covington et al., 2018). While an extensive body of literature has been devoted to alcohol’s acute aggression-heightening effects, relatively little is known about alcohol’s chronic effects. Clinical research suggests that chronic, heavy alcohol drinking can lead to a propensity for heightened aggression, including aggression perpetrated at times when alcohol was not consumed (Robertson et al., 2020; White et al., 2009). However, few preclinical studies have been devoted to studying these effects and none, to our knowledge, have examined aggression after a protracted period of abstinence.

In humans, there is substantial evidence that chronic, heavy alcohol drinking is associated with heightened aggression. Chronic, heavy alcohol intake is associated with higher scores in measures of anger, hostility, and various forms of direct and indirect aggression (Calhoun et al., 2001; Leibsohn et al., 1994; Schonwetter & Janisse, 1991; Sheehan et al., 2016), and studies have found strong associations between chronic, heavy alcohol intake and subsequent violent offending (Fergusson & Horwood, 2000; Friedman et al., 1996; Jennings et al., 2015; Komro et al., 2000; Menard & Mihalic, 2001; Orlando et al., 2005; Swahn & Donovan, 2006; Zhang et al., 1997). Furthermore, alcohol use disorder is highly comorbid with antisocial personality disorder (Helle et al., 2019), and a diagnosis of intermittent explosive disorder is associated with substance misuse (Coccaro et al., 2016). In addition, particular patterns of alcohol intake appear to have more detrimental effects with respect to aggression. Aggression and impulsivity related problems are more severe in heavy compared with moderate drinkers (Beseler et al., 2012; Siciliano et al., 2013; Townshend et al., 2014). Aggression increases with increasing severity of Alcohol Use Disorder (AUD) (Beseler et al., 2012; Kumar Sharma et al., 2012), and individuals who display a binge-like pattern of repeated intoxication and withdrawal tend to be particularly prone to aggression-related problems (Miczek et al., 2015; Waterman et al., 2019). Similarly, individuals who show binge like patterns of drinking display greater neural deficits than those who consume the same amount but in a non-binge-like pattern (Maurage et al., 2012). A repeated binge-like pattern of intoxication may have particularly deleterious consequences, not only because of the amount but also because of the specific pattern of intense use in a short period of time (Hermens et al., 2013; López-Caneda et al., 2013).

While the above-mentioned studies demonstrate that there are general associations between chronic, heavy alcohol use and aggression, they have one major shortcoming. They, for the most part, do not specify whether alcohol was consumed at the time of the aggressive encounters. This is significant because if alcohol was consumed during these encounters, then the observed associations may simply reflect the unsurprising fact that the more often someone is intoxicated, the more likely they are to experience an aggressive encounter while intoxicated. In other words, these studies are explainable in terms of acute effects alone and do not necessarily support the claim that alcohol has chronic effects. To address this, it is necessary to distinguish between *alcohol-involved*, and *alcohol-uninvolved aggression* (Robertson et al., 2020; White et al., 2009). We define alcohol-involved aggression as aggressive behavior perpetrated while the individual is acutely intoxicated. Alcohol-uninvolved aggression refers to aggressive encounters in which the perpetrator has a history of alcohol use but is in abstinence or at least is not presently intoxicated. There are, then, two kinds of chronic effects that alcohol may have. (1) Chronic alcohol intake may lead to heightened aggression in alcohol-uninvolved incidents, or (2) chronic alcohol use may cause a sensitization to alcohol’s acute effects, so that individuals with a history of heavy use respond more aggressively while intoxicated than those without such a history. To study either of these effects, it is necessary to specify whether alcohol was consumed at the time of the behavior.

In support of (1), there is mounting evidence that chronic alcohol intake is a risk factor for *alcohol-uninvolved* incidents of aggression. A study by Robertson et al. (2020) found that chronic alcohol intake was associated with aggression in college students, including aggression perpetrated at times when alcohol was not consumed. Ziherl et al. (2007) found that recovered patients with Severe Alcohol Use Disorder (SAUD) (3 years abstinent) showed higher levels of indirect aggression, irritability, negativism, suspicion, and resentment compared with non-alcoholics. Another study found that a diagnosis of SAUD within the past three years, rather than current alcohol intake, was related to the number of fights or intimate partner violence (IPV) (Leonard, 1984). Additionally, anger, hostility, and impulsiveness have been identified as features of alcohol withdrawal and may be related to withdrawal intensity (Butryn & Zeichner, 1997; Moeller & Dougherty, 2001; Ozsoy et al., 2006; Ozsoy & Esel, 2008; Shen et al., 2024). Furthermore, the Cambridge Study in Delinquent Development, found significant associations between problem drinking and subsequent violent offending, and these relationships held up when problem drinking typologies were restricted to ages 18-32 while violent offending was restricted to ages 40-50. (Jennings et al., 2015). It has also been demonstrated that abstinent alcoholics show an exaggerated hostility bias (the tendency to perceive neutral stimuli as threatening) as well as hostile attribution bias (the tendency to perceive negative outcomes as the result of others’ hostility) (Dethier & Blairy, 2012; Freeman et al., 2018; Hoffman et al., 2019; Maurage et al., 2008, 2009; Philippot et al., 1999). Finally, abstinent individuals with a history of aggressive behavior respond more aggressively in a Point Subtraction Paradigm (PSAP) in proportion to their history of alcohol intake (Ritter et al., 2019). In support of (2), several studies have shown that a history of heavy episodic drinking is a risk factor for *alcohol-involved* aggression (Kingree & Thompson, 2015; Wells et al., 2005; White & Hansell, 1996). However, these fall into the same pitfall that was previously described. That is, it is not clear whether these associations are due to a pharmacological effect of repeated alcohol intake, or whether the greater frequency of alcohol-involved aggressive encounters simply reflects a greater frequency of intoxication.

There is also limited preclinical evidence in support of both (1) and (2). In support of (1), Hwa et al. (2015) found evidence that repeated cycles of alcohol intoxication (24 h 2 bottle choice, 3 days per week) are associated with an increased likelihood of engaging in aggressive behaviors 6-8 hours after alcohol access (i.e. alcohol-uninvolved aggression). However, changes in fighting performance were not observed. In support of (2), Covington et al. (2018) found that repeated alcohol administrations, via oral gavage, at higher doses (1.8-2.2 or 2.4 g/kg) heightens the motivation to engage in a fight, measured using a Fixed Interval (FI) schedule of reinforcement. Alcohol challenge, however, did not produce changes in fighting performance. Importantly, the first administration of alcohol tended to suppress aggressive responding, whereas repeated administrations produced a sensitization of responding. Sensitized rates of FI responding for aggression were observed over a month after alcohol pretreatment.

While these clinical and preclinical studies support the claim that chronic alcohol intake can produce long-term changes in aggression, there are multiple ways to explain this association in humans. First, a winner effect may arise from past experiences of fighting while intoxicated (Hsu et al., 2006). Second, there is substantial evidence that prior aggressivity may predict subsequent alcohol intake (Airagnes et al., 2017; Chester & DeWall, 2018; Farrington, 1995; Robins, 1970; Skara et al., 2008; White et al., 1993; White & Hansell, 1996), and that this may explain apparent differences in aggression observed during abstinence (Moeller & Dougherty, 2001; Ozsoy et al., 2006). Third, a common cause, such as trait disinhibition or impulsivity may predict both alcohol use and aggression (Iacono et al., 1999; Kendler et al., 2003; Krueger et al., 2005, 2009; Latzman & Vaidya, 2013; Zucker et al., 2011). Fourth, the relationship between chronic alcohol intake and aggression may be explained by complex non-psychopharmacological factors, such as sociodemographic variables, the context of substance use, or systemic factors (Collins, 1993; Goldstein, 1985; Hoaken et al., 2012; Lipsey et al., 1997). Finally, the hypothesis which we propose here is that chronic alcohol intake may directly cause long term changes in aggressive behavior through psychopharmacological mechanisms. It is likely, however, that a combination of these factors may explain the observed relationships, and that the various causal pathways may feed into each other. For example, studies have found that while prior aggression predicts subsequent heavy drinking, heavy drinking, in turn, predicts heightened aggression even after controlling for prior aggression and other variables (Hussong et al., 2001; Najman et al., 2019).

While multiple of the above explanations may be valid and operative in the relationship between chronic alcohol intake and aggression, we hypothesize that chronic alcohol intake can directly cause long term changes in aggression. The goal of the present study was to examine whether chronic binge-like alcohol drinking produces changes in either alcohol-uninvolved or alcohol-involved aggressive behavior. We employed a highly translational model of binge-like drinking, Drinking in the Dark (DID) and examined aggression during acute and protracted abstinence. DID has traditionally been used in high-drinking C57BL/6J mice and has been shown to generate significant changes in behavior after a period of abstinence (Neira et al., 2022; Thiele & Navarro, 2014). We applied this model to genetically heterogeneous CFW mice in order to capture individual differences and subpopulations in both drinking and aggressive behaviors. We aimed to examine the trajectory of these behaviors within subjects over the course of repeated cycles of binge drinking. We used a neutral-arena aggression paradigm to sensitively detect bidirectional changes in aggression (Fish et al., 1999; Miczek & O’Donnell, 1978; Newman et al., 2015). We also sought to investigate whether a single session of binge-like drinking causes acutely heightened aggression, and whether this effect is dependent on prior chronic drinking histories.

We found that repeated cycles of binge-like drinking heightens aggression in alcohol-uninvolved fights. This effect occurs in heavy drinkers but not in moderate drinkers given 5-8 weeks of alcohol access and approximately 20 hours of abstinence. After 9 weeks of DID and one full week of abstinence, we found that animals with a history of binge-like alcohol drinking initiate a fight more rapidly and with more consistency than H₂O controls. Finally, we found that a single session of voluntary binge-like drinking, promoted by water restriction and SuperSac vehicle, produces heightened aggression in both alcohol-naïve animals and animals with substantial alcohol histories, and the magnitude of heightened aggression is directly correlated with the animals’ blood ethanol concentration (BEC) after the fight. This study presents a novel model for examining causal sequences in the relationships between repeated binge-like drinking and aggression which are difficult to disentangle in human research.

## Materials and Methods

### Subjects

Eleven-week-old male Swiss Webster (CFW) mice (Charles River Laboratories, Raleigh, NC, USA) were pair housed with age matched, ovariectomized CFW females for at least 2 weeks before aggressive encounters to facilitate territorial agonistic behavior toward male conspecifics (Miczek & O’Donnell, 1978). All mice were housed in clear polycarbonate cages (30 × 19 × 14 cm) lined with pine shavings, and had unlimited access to tap water and rodent chow (PicoLab Laboratory Rodent Diet 5L0D) through stainless steel wire mesh cage lids. Food, water, and bedding were changed weekly at least 24 h prior to behavioral testing. The vivarium was temperature-regulated and maintained on a 12-h reverse light/dark photocycle (lights off from 0700 to 1900 h). Animals were cared for according to the National Research Council Guide for the Care and Use of Laboratory Animals, and procedures were approved by the Tufts University Institutional Animal Care and Use Committee.

### Baseline Behavioral Screening

#### Home cage screening

After at least two weeks of pair bonding, 21 CFW males were screened for baseline aggression in their home cage using a resident-intruder paradigm previously described (Miczek & O’Donnell, 1980; Newman et al., 2015). Home cage screening occurred for at least 5 days, such that residents displayed consistent aggression towards an intruder male. Females were removed from the resident’s home cage before an intruder male was introduced into the cage. Intruders were age-matched, group housed male C57BL/6J mice. Intruders were regularly cycled to prevent resident-intruder habituation, and aggressive intruders were excluded from further testing. Aggressive encounters were terminated 5 min after the resident initiated the first attack bite or at 5 min if the resident failed to attack. The latency to the first bite, and the bite frequencies were recorded with handheld timers and tally counters.

#### Neutral arena screening

Compared to aggression in the home cage, a neutral arena reduces aggressive behavior by approximately 50%, allowing for sensitive detection of bidirectional changes in aggressive behavior (Fish et al., 1999; Miczek & O’Donnell, 1980). After consistent aggression was established, residents were singly housed. Residents were then screened daily for one additional week with daily resident-intruder fights alternating between their home cage and a neutral arena as previously described (Miczek & O’Donnell, 1978; Newman et al., 2015). The neutral arena was a large (48 x 26 x16 cm) clean polycarbonate cage lined with fresh pine shaving bedding. Before residents were introduced into the neutral arena, they were screened for baseline locomotor activity in an empty plexiglass cage of the same dimensions to ensure that there were no differences in baseline locomotor response to the novel environment (Fig. S1). Mice were placed in one corner of the arena and a plexiglass lid was placed on top. Locomotor activity was recorded for 5 minutes using Noldus Ethovision XT with an overhead camera (Panasonic WV-CP280). In neutral arena fights, residents were dropped into one corner of the neutral arena and the intruder was introduced in the opposite corner after a 5 second delay. Aggressive encounters were terminated 5 min after the resident initiated the first attack bite or at 5 min if the resident failed to attack. The cage was cleaned between each fight to eliminate pheromonal cues.

Residents were then assigned to either H₂O (n = 8) or EtOH (n = 12) drinking groups, counterbalanced by their average bite frequencies in the last two neutral arena fights. One animal was excluded from the study after displaying consistent non-aggression towards an intruder. Before proceeding to Drinking in the Dark, we ensured that there were no significant differences in bite frequency or latency either in the home cage or the neutral arena (Fig. S2), and that there were no differences in locomotor activity (Fig. S1). We also confirmed that neutral arena aggression was approximately 50% that of the home cage in terms of both bite frequency and latency for both groups. To control for potential order effects, behavioral testing alternated between EtOH and H₂O drinkers.

### Voluntary Binge-Like alcohol intake: Drinking in the Dark

Mice were given access to either 20% EtOH (w/v) (n = 12) or H₂O (n = 8) following a Drinking in the Dark (DID) protocol, as described previously (Thiele & Navarro, 2014). Fluid access was provided four days a week, starting 3 h into the dark cycle. Animals were given 2 h of access on Mondays, Tuesdays, and Wednesdays, and 4 h of access on Thursdays. Animals were 20 weeks of age at the start of alcohol access and DID was conducted for 9 consecutive weeks. Fluid was dispensed via modified 10ml serological pipets fixed to double ball bearing, stainless steel sippers (Braintree Scientific). The sipper tubes were inserted into the cage through wire mesh lids and protruded into the cage approximately the same distance as their normal water bottle sippers (∼4 cm). The tubes were fixed to the cage lid with binder clips to prevent leakage induced by animal movement. Intake was measured by recording fluid volumes at the beginning of the session, as well as at 20 min and 2 h time points. On 4 h days (i.e., Thursdays), fluid volumes were recorded at 20 min, 2 h and 4 h time points. To control for normal leakage due to dripping, two empty ‘drip cages’ were maintained during all drinking sessions, one of which contained an EtOH filled tube, and the other contained an H₂O tube. Drip values were subtracted from all the intake data of that day. All drinking tubes were sanitized daily by soaking in Alconox solution and thorough rinsing with tap water. The binder clips were kept separate for each animal to ensure that there was no cross contamination of pheromonal cues.

### Resident-intruder Aggression Tests During Acute and Protracted Abstinence

Acute abstinence aggression testing occurred on Fridays at least 20 h after the end of the drinking session on the previous day. Resident-intruder tests were conducted in the neutral arena after 1, 4, 5, 6 and 8 weeks of DID using the same protocol described above. Intruders were 11-week-old, novel male C57BL6J mice with a moderate history of defeat (approximately 2-4 previous defeat experiences). After 9 weeks of DID, mice underwent 1 week of abstinence with no behavioral testing, followed by four consecutive days of aggression testing, with resident-intruder tests alternating between the home cage and the neutral arena (Fig. 1A).

**Fig 1.**
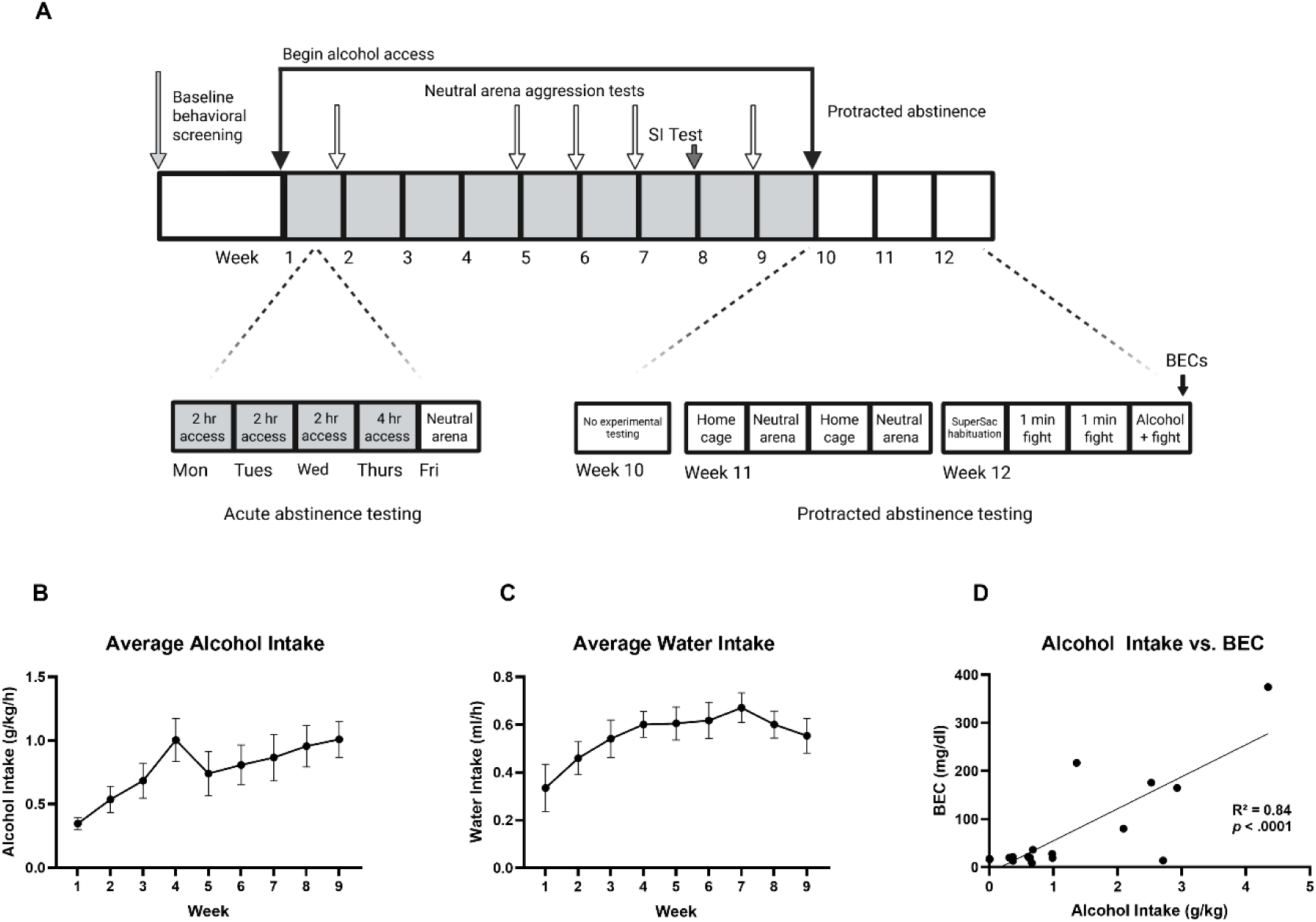
Experimental timeline and intake measures. After baseline behavioral screening, mice underwent 9 weeks of Drinking in the Dark (DID) with access to either 20% EtOH (n = 11-12) or H₂O (n = 8). Alcohol or water access was provided for 2 h on Mondays through Wednesdays and 4 h on Thursdays. Resident-intruder aggression tests were conducted in a neutral arena on Fridays (∼20 h post-drinking) in weeks 1, 4, 5, 6, and 8. A social interaction (SI) test was conducted in Week 7. Further aggression testing was conducted during protracted abstinence in week 11. In week 12, all animals underwent a single session of voluntary binge-like alcohol intake followed by a resident intruder test. BECs (mg/dl) were collected 30 minutes following this test **(A)**. Average weekly alcohol intake (g/kg/h) increased over the first 4 weeks and stabilized between Weeks 4–9 (RM ANOVA: F (2.63, 26.27) = 6.55, p = .003) **(B)**. Water drinkers showed a similar trend in water intake (ml/h) (F (3, 21) = 3.82, p = 0.025) **(C)**. Alcohol intake (15% EtOH in SuperSac, reported in g/kg) during the final 1 h drinking session was strongly correlated with BECs collected 30 min post-drinking (R² = 0.8439, *p* < 0.0001) **(D)**. Data are represented as mean ± SEM. Experimental timeline graphics were created with Biorender.com.

### Social Interaction Test

After 7 weeks of DID and 20 hours of abstinence, mice underwent a social interaction test using an apparatus described previously (Berton et al., 2006; Golden et al., 2011) (Fig. 1A). The social interaction arena was a 42 cm X 42 cm open field custom-crafted with opaque black Polyvinyl Chloride. A wire-mesh shark cage secured in clear Plexiglas was centered against one wall of the arena for presentation of a stimulus animal (Fig. 2A). Each resident was placed in one corner of the arena for a 2.5 minute ‘non interaction test’ in which no stimulus animal was present. The resident was then placed back in their home cage while a stimulus animal was placed into the wire mesh enclosure. The resident was then placed back into the arena for an additional 2.5 minute ‘interaction’ test. The interval between non-interaction and interaction tests was no more than 30 s. The sessions were recorded with an overhead camera in dim red light, and the videos were processed with Noldus Ethovision XT. Locomotor activity was recorded, as well as time spent in the ‘interaction zone’ (a 14 cm X 24 cm rectangle projecting 8 cm around the wire mesh enclosure) and in the ‘corner zones’ (a 9 cm X 9 cm area projecting from each corner joint opposite the interaction zone). The stimulus animals were non-threatening male CFW mice. To control for differences in testosterone and aggressive behavior, all stimulus animals were castrated and screened for non-aggression towards an intruder prior to social interaction testing.

**Fig 2.**
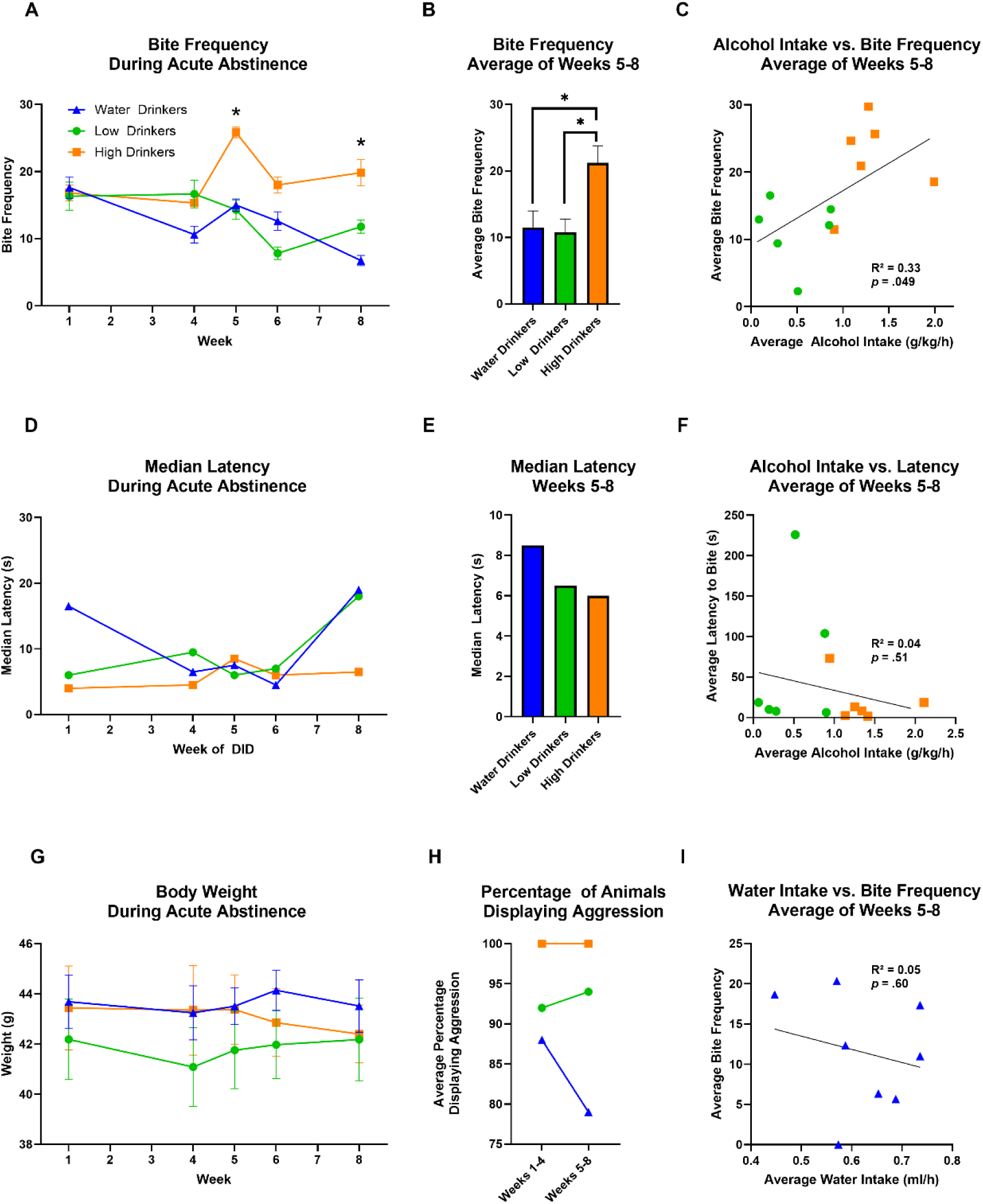
High drinkers bite more frequently in neutral arena fights following 5-8 weeks of DID. Bite frequency was measured in Water Drinkers (n = 8), Low Drinkers (n = 5-6), and High Drinkers (n = 6) during neutral arena fights following 1, 4, 5, 6, and 8 weeks of DID and 20 h of abstinence. A significant effect of drinking group was observed at Weeks 5 and 8 (Week 5: H(2) = 8.30, *p* = .016; Week 8: H(2) = 6.39, *p* = .041) **(A)**. When averaged across Weeks 5, 6, and 8, High Drinkers exhibited significantly greater bite frequencies than both Water (*p* = .028) and Low Drinkers (*p* = .029), with no difference between Water and Low Drinkers (ANOVA: F(2,17) = 5.31, *p* = .016) **(B)**. Alcohol intake positively correlated with bite frequency over this period (R² = 0.33, *p* = .049) **(C)**. Median latency to bite (s) did not differ across groups at any time point (*p* > .05) **(D)**, nor when the median latency was calculated for all fights across Weeks 5–8 (H(2) = 1.92, *p* = .38) **(E)**. Alcohol intake was not significantly associated with latency (R² = 0.04, *p* = .51) **(F)**. High Drinkers maintained 100% aggression rates from Weeks 1 to 8, while Water Drinkers showed a decline in aggression rates between early DID (weeks 1-4) and late DID (weeks 5-8) **(G)**. No differences in body weight (g) were observed between groups during DID (F(1, 6) = 0.45, *p* = .529) **(H)**. Among Water Drinkers, average water intake (ml/h) over weeks 5-8 was not significantly associated with average bite frequency (R² = 0.05, *p* = .60) **(I)**. Data are shown as mean ± SEM. Latencies are represented as median values. Asterisks indicate statistically significant differences (*p* < .05).

### Acute Voluntary Binge-Like Alcohol Drinking Prior to Fight

After 3 weeks of abstinence, mice underwent a voluntary alcohol drinking test (Fig. 1A). All mice were habituated to consume SuperSac vehicle (0.125% saccharine *w/v*, 4% glucose *w/v* in tap water) (Valenstein et al., 1967). Mice were water restricted for 1.5 h, then given 30 min of access to the SuperSac vehicle. This was repeated until all animals consumed at least 0.5 ml of SuperSac within a single 30 min session. All mice met the criteria for supersac intake after 3 sessions of access. Mice then underwent two days of home cage aggression screening with the fight duration reduced to 1 min. The following day, mice were water restricted for 1 h, then given 1 h of access to 15% EtOH in SuperSac vehicle using the same tubes in which they had previously consumed 20% EtOH or H_2_O during DID. Intake was also measured using the same methods: recording pre and post volumes and subtracting drip values. After tubes were removed, animals were immediately presented with an intruder in their home cage and a 1 min fight ensued. Animals were sacrificed 30 minutes after the fight. Trunk blood was collected in heparinized tubes, spun down, and serum was analyzed for Blood Ethanol Concentration with the EnzyChrome ECET Ethanol Assay Kit.

### Statistics

All analyses were conducted with α set at 0.05 using SPSS Statistics (IBM Corp., Armonk, NY). Data were examined for normality prior to statistical testing. When assumptions of normality or equal variance were violated, nonparametric tests were applied as appropriate. These included Friedman’s test for within-subject comparisons, Kruskal–Wallis tests for group comparisons involving three or more conditions, and Fisher’s exact test for categorical variables with small sample sizes. For resident-intruder tests, animals that failed to initiate an attack were assigned the maximum possible latency, and latency data were analyzed using nonparametric tests. Median values are reported in all latency figures except for XY plots. When linear regressions were conducted with latency data, animals that failed to attack were excluded. Alcohol intake and other continuous, normally distributed variables were analyzed using one-way or repeated measures ANOVAs. Significant main effects or interactions were followed by appropriate post hoc comparisons. Graphs and figures were generated in GraphPad Prism (version 10.4.1; GraphPad Software, San Diego, CA) with all data represented as mean ± SEM unless otherwise noted.

## Results

### CFW mice show a range of alcohol intake in Drinking in the Dark

Alcohol intake in CFW mice increased over the first 4 weeks of DID, beginning at an average of 0.35g/kg/h in the first week, and stabilizing at an average of 0.93 g/kg/h between weeks 5-9 (Fig. 1B). A repeated measures ANOVA using Greenhouse-Geisser correction revealed a significant effect of time on average alcohol intake over the course of weeks 1-9 of DID (F (2.63, 26.27) = 6.55, *p =* .003). No significant effect of time was found between weeks 4-9 alone (F (5,50) = 2.01, *p* > .05), nor between weeks 5-9 alone (F (4,40) = 2.11, *p* > .05), indicating that alcohol intake stabilized over the latter half of DID. Mice in the water group also showed an escalation in water intake over the first 4 weeks, beginning at an average of 0.34 ml/h, and stabilizing at an average of 0.61 ml/h over weeks 5-9 (Fig. 1C). Because of the smaller sample size of the water group, water intake was compiled into 2 week bins spanning weeks 1-8 before a repeated measures ANOVA was conducted, revealing a significant effect of time (F (3, 21) = 3.82, *p = .*025). Alcohol intake tended to escalate over the week (Monday through Thursday drinking sessions), with the lowest levels of intake on Mondays (approximately 75% of the weekly average) (Fig. S3). A Friedman test was revealed a statistically significant difference in alcohol intake across weekdays (**χ²** (3) = 10.30, *p = .*016). Median ranks increased from Monday (1.50) to Thursday (3.08), suggesting intake tended to rise throughout the week. Water intake, similarly, tended to increase from Monday to Wednesday, but also showed a decrease on Thursdays (Fig. S4). A repeated measures ANOVA confirmed that there was a significant effect of weekday on water intake (F (3, 21) = 4.11, *p = .*019), and trend analysis further indicated that water intake followed a cubic pattern (F(1, 7) = 8.22, *p =* .024).

Although alcohol intake was stable within subjects over weeks 5-9, there was considerable variability between subjects, with intake ranging from 0.12 g/kg/h to 1.92 g/kg/h. Because of these individual differences in alcohol intake in CFW mice, we classified animals into high and low drinking groups via median split (median = 1.04g/kg/h, ‘High Drinkers’ mean = 1.33 g/kg/h, ‘Low Drinkers’ mean = 0.45 g/kg/h). Baseline measures of aggression, obtained from an average of the last two screening sessions, were not predictive of subsequent high or low drinker status (neutral arena latency, t (10) = 1.34, *p = .*24, neutral arena bite frequency, t (10) = −0.95, *p = .*36, home cage latency, t (10) = 1.37, *p = .*20, home cage bite frequency, t (20) = −0.32, *p = .*75). Frontloading was assessed by comparing the rate of intake (g/kg/min) during the first 20 minutes of each drinking session with the rate of intake in the remainder of the session. Rates were averaged across weeks 5–9 to calculate a “rate ratio” (early vs. late session intake). Animals with a rate ratio greater than 1 were classified as frontloaders. Similarly, Water Drinkers were classified as frontloaders based on positive early-to-late rates of water intake rates in ml/min. 50% of High Drinkers, 17% of Low Drinkers showed frontloading behavior (Fig. S5). Additionally, 50% of Water Drinkers frontloaded water. However, alcohol frontloaders exhibited an 87% increase in intake rate during the initial 20 minutes relative to the remainder of the session, whereas water frontloaders showed a 55% increase in early intake rate. BECs were measured following the acute alcohol drinking and aggression test, after all animals had consumed 15% EtOH in SuperSac using the same tubes in which they had previously consumed water or 20% EtOH during DID. We verified that there was a significant correlation between alcohol intake measured by volume and BECs using the ECET ethanol plate assay (R² = 0.8439, p < 0.0001) (Fig. 1D). From these data, we estimate that High Drinkers achieved BECs of approximately 70-90 mg/dl/h during weeks 5-9 of DID, while Low Drinkers achieved BECs of approximately 10-20 mg/dl/h.

### High Drinkers become more aggressive following 5-8 weeks of drinking in the dark

After 1 week of DID, there were no differences in resident-intruder aggression between High Drinkers, Low Drinkers, and H₂O control animals when testing was performed during acute abstinence ∼24 h after the last DID session (i.e., ‘alcohol-uninvolved’ aggressive behavior). Over the course of repeated DID cycles, however, group differences began to emerge. Both H₂O drinkers and Low Drinkers showed a gradual decline in bite frequency over the course of DID while High Drinkers maintained bite frequencies at or above baseline levels over the course of weeks 5 - 8 (Fig. 2A). Data were not normally distributed at some time points. Therefore, we conducted a series of Kruskal-Wallis tests, which revealed significant effects of drinking group at week 5 (*H*(2) = 8.30, *p =* .016), and at week 8 (*H*(2) = 6.39, *p =* .041). No significant group differences were found at weeks 1 (*p =* .918), 4 (*p =* .213), or 6 (*p =* .095). In order to further probe the observed group differences between weeks 5 and 8, we calculated the average bite frequency data over weeks 5,6, and 8, which resulted in a normally distributed dataset on which a One Way ANOVA could be performed. This revealed a significant effect of drinking group on bite frequency (F(2, 17) = 5.31, *p =* .016). Post hoc comparisons using the Tukey HSD test revealed that High Drinkers exhibited a significantly higher average bite frequency compared to both Water Drinkers (*p =* .028) and Low Drinkers (*p =* .029), but there was no significant difference between bite frequencies of Water Drinkers and Low Drinkers (*p* > .05). High Drinkers’ average bite frequency was approximately double that of both Low Drinkers and Water Drinkers during this time (Fig. 2B). Additionally, there was a significant positive correlation between alcohol intake and bite frequency when data was averaged over weeks 5 - 8 (R² = 0.3348, *p = .*049) (Fig. 2C).

Latency during acute abstinence was variable and deviated from normality. A series of Kruskal–Wallis tests revealed no significant differences in bite latency across drinking groups at any time point during the 8-week drinking period. Specifically, there were no group effects observed at week 1 (*H*(2) = 1.12, *p =* .57), week 4 (*H*(2) = 0.62, *p =* .73), week 5 (*H*(2) = 0.50, *p =* .78), week 6 (*H*(2) = 0.08, *p =* .96) or week 8 (*H*(2) = 1.86, *p =* .40) (Fig. 2D). Similarly, no significant effect of drinking group was observed when data was compiled over weeks 5-8 (*H*(2) = 1.92, *p =* .38) (Fig. 2E), nor was there a significant correlation between alcohol consumed over weeks 5-8 and latency to bite during that time (R² = 0.043, *p =* .51) (Fig. 2F). We noted, however, that the median latency over weeks 5-8 was lower in High Drinkers compared with the other groups by a small margin (Fig. 2E), and that High Drinkers were generally less susceptible to fluctuations in latency (Fig. 2D). High Drinkers also displayed more consistent aggression compared with Low Drinkers or Water Drinkers and were the only group to maintain 100% aggression rates at all time points (Fig. 2H). This is reflected in their consistently low median latencies.

In order to ensure that the observed changes in aggressive behaviors were not due to general differences in vitality, viability, or caloric intake between groups, we verified that these changes in aggression were not explained by differences in baseline locomotor activity (collected prior to the start of DID), nor by differences or trends in body weight. High or Low Drinker status was not related to baseline locomotor activity in terms of either velocity (*t*(6.13) = –0.57, *p =* .587) or distance traveled (*t*(6.13) = –0.57, *p =* .587). Additionally, residents’ body weights were measured prior to each fight across the duration of DID. A repeated measures ANOVA revealed no significant effect of time on body weight (F(4, 24) = 1.74, *p =* .174), no significant interaction between time and drinking group (F(4, 24) = 1.21, *p =* .332), and no significant difference in body weight between groups (F(1, 6) = 0.45, *p =* .529) (Fig. 2G), further suggesting that the observed behavioral changes were not due to differences in caloric intake or viability. Additionally, we observed no significant association between water intake and bite frequency over weeks 5-8 (R² = 0.05, *p =* .60). In contrast to the high alcohol drinking group, high water drinkers showed modestly lower aggression than low drinkers (Fig. 2I), suggesting that elevated bite frequency in High Drinkers is not simply reflective of a general relationship between consummatory behaviors and aggression.

### High Drinkers, Low Drinkers, and Water Drinkers show no differences in locomotor activity or social interaction during acute abstinence

To further verify that the observed differences in aggression were not merely an artifact of general differences in viability, we conducted a social interaction test at week 7 of DID. The test involved 2.5 mins of free exploration in an open field, followed by 2.5 mins in the same arena with a stimulus animal (castrated male CFW mouse) presented in a shark cage (Fig. 3A). In the initial non-interaction portion of the test, a one-way ANOVA revealed no significant differences in locomotion between High Drinkers, Low Drinkers, and Water Drinkers in terms of distance traveled (F(2, 16) = 0.93, *p =* .41), or velocity (F(2, 16) = 0.93, *p =* .41) (Fig. 3B and Fig. 3C). There were also no significant group differences in the interaction ratio (the time spent in the interaction zone during the interaction vs. the no-interaction tests) (F(2, 16) = 0.93, *p =* .41) (Fig. 3D), nor in the corner ratio (proportion of time spent in the corners during the interaction vs. the no-interaction tests) (F(2, 16) = 1.42, *p =* .27) (Fig. 3E). We noted, additionally, that High Drinkers displayed moderately higher levels of social interaction and spent moderately less time in the corners in the presence of a stimulus animal, but no similar trend in locomotor activity. In short, despite substantial differences in aggressive behaviors, all three groups maintain similar levels of locomotor activity (both in the presence and in the absence of another animal) and show no differences in sociability (either in approach or avoidance behaviors). This suggests that the observed differences in both drinking and aggression are not likely to be artifacts of general differences in viability.

**Fig 3.**
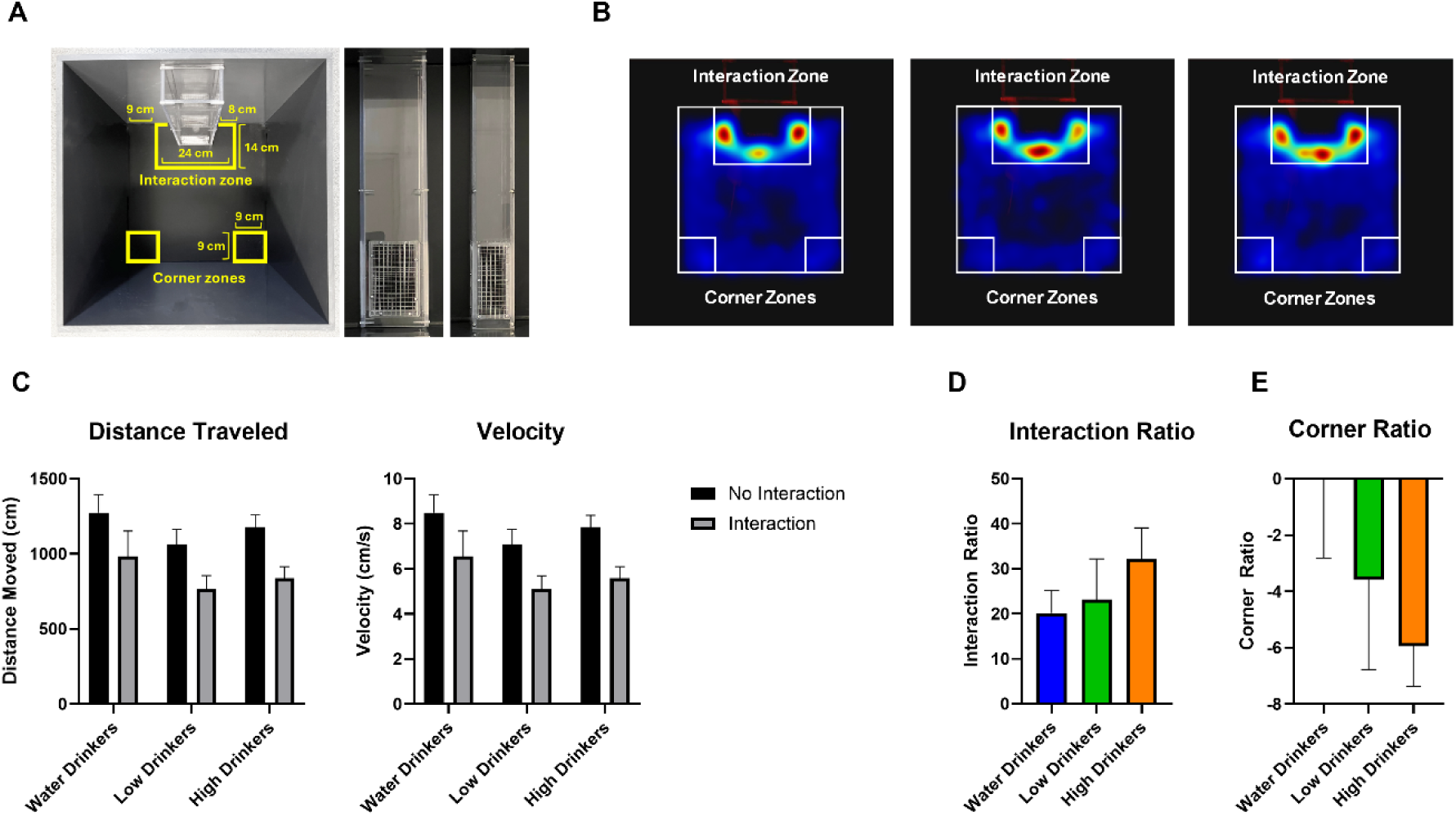
A social interaction test was conducted after 7 weeks of DID and 20h of abstinence. Water Drinkers (n = 8), Low Drinkers (n = 5), and High Drinkers (n = 6) underwent a 2.5 min no-interaction trial followed by a 2.5 min interaction trial with a castrated male stimulus animal placed in a Plexiglas shark cage. Zones were defined as shown: a central interaction zone was defined surrounding the shark cage and the opposite corners were defined by corner zones **(A)**. Representative heatmaps show similar spatial occupancy patterns across groups, represented as composites of animal center-point locations across all trials **(B)**. No significant group differences were found in total distance traveled (cm) (F(2, 16) = 0.93, *p* = .413) or velocity (cm/s) (F(2, 16) = 0.93, *p* = .41) across both no-interaction and interaction trials **(C)**. Interaction ratios (time spent in the interaction zone during interaction vs. no-interaction trials) did not differ between groups (F(2, 16) = 0.93, *p* = .41) **(D)**. Corner ratios (time spent in corner zones during interaction vs. no-interaction trials) also did not differ between groups (F(2, 16) = 1.42, *p* = .27) **(E)**. Data are shown as mean ± SEM.

### In the first aggressive encounter following 1 week of abstinence, chronic alcohol drinkers were faster to initiate a fight

After 9 weeks of DID, mice underwent one week of abstinence with no behavioral testing followed by four days of resident-intruder tests, alternating between the home cage and the neutral arena (Fig 1A). In the first resident-intruder test, conducted in the home cage, Water Drinkers exhibited significantly greater variability in bite latency compared to both High and Low Drinkers (Levene’s test: *F*(2, 14) = 22.74, *p* < .001). Additionally, latency values within the Water group deviated from normality (*Shapiro–Wilk* = .750, *p =* .020) and appeared bimodally distributed, with approximately half the animals displaying markedly prolonged latencies to bite, while the others maintained short latencies (Figure 4B). To address the small sample size and non-normal distribution of the data, we examined the proportion of animals that had initiated a fight over the course of the 5 minute encounter in 15 s increments (Fig 4B). We compared Water Drinkers to all Alcohol Drinkers using a series of Fisher’s Exact Tests, which revealed that a significant group difference emerged after 30 s (*p =* .011). Specifically, by this time point, 100% of alcohol drinkers had initiated a fight, while less than 50% of Water Drinkers had done so (Fig. 4B and Fig. S6). This group difference remained significant between 45 s (*p =* .043) and 150 s (*p =* .043), but was no longer significant at later time points (e.g., *p =* .197 at 165 s), suggesting that chronic alcohol drinkers are quicker to initiate a fight during the initial aggressive encounter following a period of protracted abstinence, compared with alcohol naïve animals. It is also notable that while there were no significant group differences in body weight during protracted abstinence (*F*(2, 14) = 0.85, *p* = .45), Water Drinkers gained relatively more weight than alcohol drinkers after the end of DID and remained slightly heavier than alcohol drinkers throughout protracted abstinence (Fig. 4D). This suggests that Alcohol Drinkers’ quick latencies to initiate the first fight was not due to greater vitality or bodyweight compared with Water Drinkers. After repeated daily testing, Water Drinkers’ bite latencies dropped to levels similar to the alcohol groups. This was expected, as bite latencies typically decrease with repeated daily testing. We observed no significant differences in bite frequency between groups across any of the protracted abstinence test sessions. Due to non-normality in portions of the data, we conducted Kruskal–Wallis tests, which confirmed no significant group effects in Session 1 (H(2) = 0.48, *p =* .789), Session 2 (H(2) = 1.16, *p =* .559), Session 3 (H(2) = 3.87, *p =* .145), or Session 4 (H(2) = 0.23, *p =* .893). (Fig. 4C).

**Fig 4.**
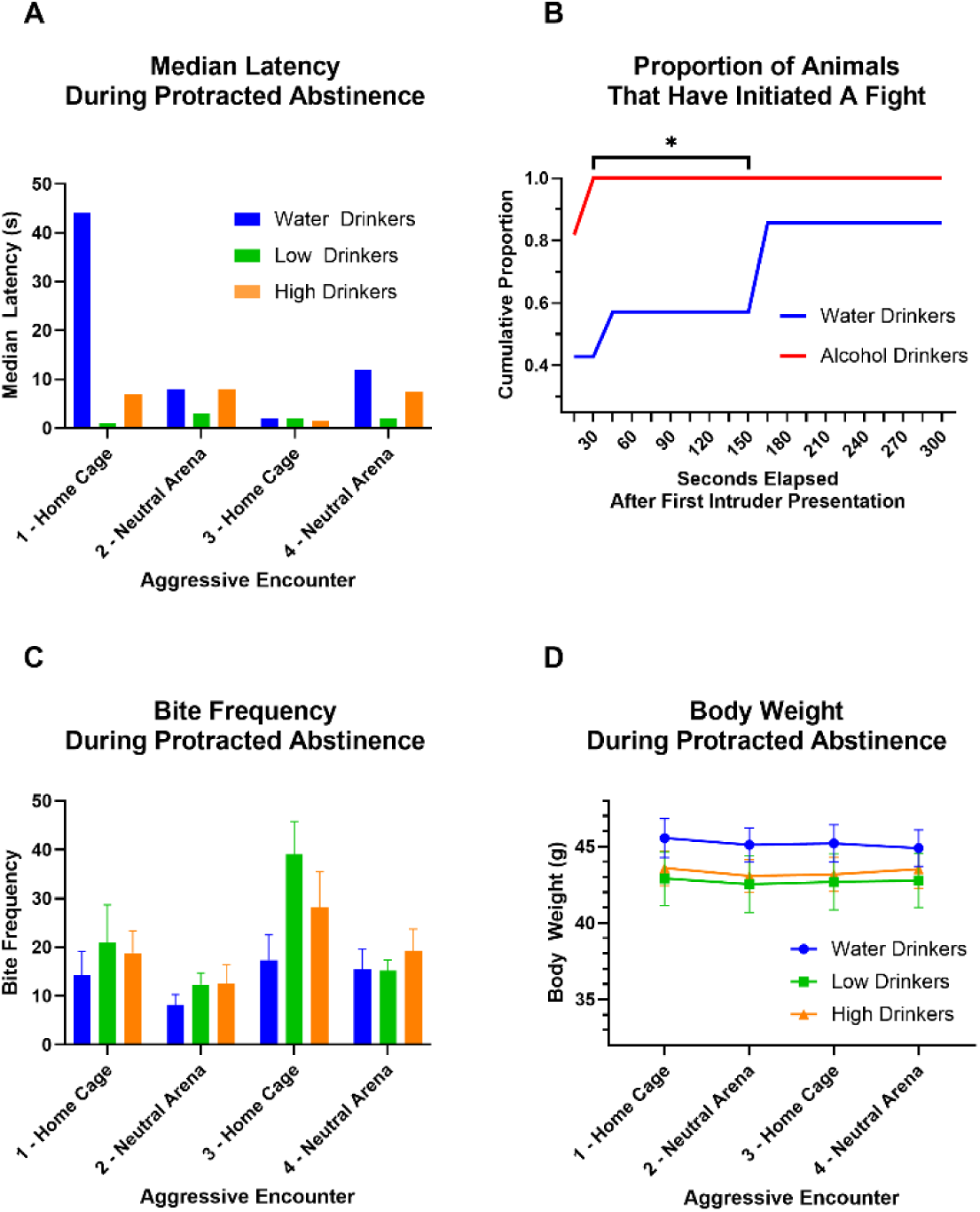
After 9 weeks of DID, animals underwent 1 week of abstinence with no behavioral testing followed by four aggressive encounters, alternating between the home cage and a neutral arena. Median latency to bite (s) is represented for Water Drinkers (n = 7), Low Drinkers (n = 5), and High Drinkers (n = 6) **(A)**. During the first aggressive encounter, alcohol drinkers were faster to initiate fights than water drinkers, as shown by the cumulative proportion of animals that initiated a fight within successive 15 s bins. All alcohol drinkers initiated a fight within 30 seconds of the first intruder presentation, whereas less than 50% of water drinkers did so. A series of Fishers Exact tests over each successive 15s time bin revealed that a significant group difference emerged at 30 s (*p* = .011) and this difference remained significant between 45 s (*p* = .043) and 150 s (*p* = .043) **(B)**. Bite frequency during protracted abstinence did not significantly differ between groups (*p* > .05) **(C)**. No significant differences in body weight (g) were observed between groups across test days during the abstinence period (*F*(2, 14) = 0.85, *p* = .45) **(D)**. Data are shown as mean ± SEM. Latencies are represented as median values. Asterisks indicate statistically significant differences (*p* < .05).

### Binge-like alcohol drinking acutely heightens aggression regardless of drinking history

We conducted a test for acute alcohol-heightened aggression across all three groups, formerly Water Drinkers, Low Drinkers, and High Drinkers, henceforth referred to as ‘Alcohol Naive’, ‘Chronic Low Drinkers’ and ‘Chronic High Drinkers’. Before the acute alcohol test, all animals were habituated to consume SuperSac vehicle using the same drinking tubes in which they had consumed water or alcohol during DID. All animals consumed at least 0.5ml of SuperSac vehicle within three 30 minutes sessions. A repeated measures ANOVA revealed no significant interaction between habituation session and drinking group (F(4, 30) = 2.134, *p =* .10). However, Chronic High Drinkers tended to consume moderately more SuperSac than either Chronic Low Drinkers or Alcohol Naive animals (Fig. 5A). After SuperSac vehicle habituation, animals were screened for 1 min fights on two consecutive days to obtain a pre-alcohol baseline. A One-way ANOVAs showed no significant group differences in bite frequency (Session 1: *F*(2, 15) = 0.27, *p =* .765; Session 2: *F*(2, 15) = 0.42, *p =* .665), and Kruskal–Wallis tests showed no significant group differences in latency to bite (Session 1: *H*(2) = 0.27, *p =* .872; Session 2: *H*(2) = 0.29, *p =* .867). The subsequent day, all animals were given access to 15% EtOH in SuperSac vehicle (to promote high intake) followed by the presentation of an intruder for a one minute fight. BECs were measured 30 minutes after the fight. Alcohol intake was relatively consistent across all drinking history groups (Fig. 5C and Fig. 5D). There were no significant differences in alcohol intake (*H*(2) = 0.76, *p =* .68) or BECs (*H*(2) = 0.42, *p =* .81) between groups (Fig. 5C and Fig. 5D).

**Fig 5.**
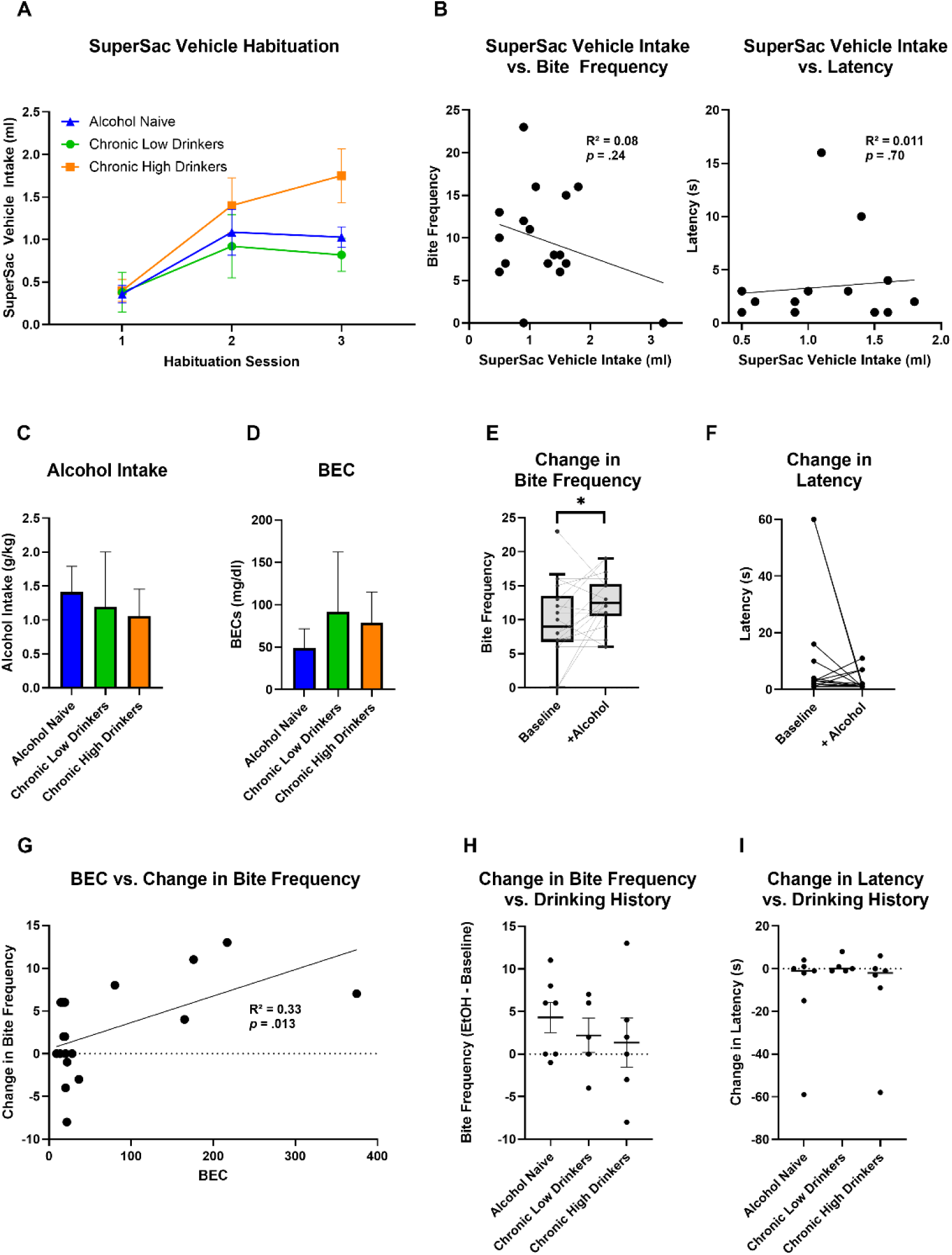
Acute binge-like alcohol intake increases aggression. Prior to testing, animals were either Alcohol Naïve (n = 7), Chronic Low Drinkers (n = 5), or Chronic High Drinkers (n = 6). Animals were habituated to consume SuperSac vehicle over the course of three 30 min habituation sessions separated by 1.5 h of water restriction. Vehicle intake (ml) increased over sessions and was moderately, but not significantly higher in Chronic High Drinkers (F(4, 30) = 2.134, *p =* .10) **(A)**. SuperSac Vehicle intake (ml) during the final habituation session was not significantly associated with either bite frequency (R^2^ = 0.084, *p = .*24) or bite latency (s) (R^2^ = 0.011, *p = .*70) in a resident intruder test conducted on the following day **(B)**. During the acute alcohol drinking test, mice were water restricted for 1 h, followed by 1 h of access to 15% EtOH in SuperSac. Alcohol intake (g/kg) did not significantly differ between groups (*H*(2) = 0.76, *p =* .68) **(C)**. BECs (mg/dl) collected 30 min after the end of the drinking session also did not significantly differ between groups (*H*(2) = 0.42, *p =* .81) **(D)**. Compared to a baseline test, a paired t test revealed a significant increase in bite frequency following acute alcohol intake (*t*(17) = –2.15, *p =* .046) **(E).** Latency to bite (s) decreased moderately but this difference was not significant (*p* > .05) **(F)**. The change in bite frequency from the baseline test was positively correlated with BECs (R^2^ = 0.33, *p = .*013) **(G)**. There were no significant group differences in the change in bite frequency from baseline (*p* > .05). However, Alcohol Naïve animals were showed the greatest increase in bite frequency **(H)**. The change in latency from baseline also did not significantly differ across groups (F(2, 19) = 1.06, *p* = .37) **(I)**. Data are shown as mean ± SEM. Latencies are represented as median values. Whiskers in box-and-whisker plots represent 10th–90th percentiles.

We found that one session of acute binge-like alcohol drinking produced heightened aggression regardless of prior drinking histories. Specifically, alcohol produced a significant elevation in bite frequency and a moderate reduction in latency to bite. A paired-samples t-test across all subjects revealed that bite frequencies were significantly higher during the fighting session that immediately followed alcohol + SuperSac intake compared to the previous day’s baseline (*t*(17) = –2.15, *p =* .046) (Fig. 5E). There was also a significant positive correlation between BECs measured after the fight and the change in bite frequency compared to the previous day’s baseline (R^2^ = 0.33, *p = .*013) (Fig 5G). This correlation remained significant for alcohol naive animals alone (R^2^ = 0.58, *p = .*047), as well for those with an alcohol drinking history (R^2^ = 0.40, *p = .*038). A one-way ANOVA revealed no significant group differences in the change in bite frequency from baseline (*F*(2, 15) = 0.49, *p =* .623) (Fig. 5H). These findings suggest that acute binge-like alcohol drinking produces escalated aggression in direct proportion with BECs at the time of the encounter. Importantly, this occurs regardless of whether it is the animal’s first exposure to alcohol, or whether they have had substantial experience with alcohol drinking.

While bite frequency was significantly elevated across groups, latency was moderately but not significantly reduced following the alcohol drinking session. A Friedman test revealed no significant change in latency between baseline and the post-alcohol fight (χ²(1, N = 18) = 1.667, *p =* .197). However, we observed a moderate reduction in latency, mirroring the observed escalation in bite frequency (Fig. 5F). Similarly, bite latencies were not significantly associated with BECs (R^2^ = 0.015, *p = .*65) (Fig S7), and a Kruskal-Wallis test revealed no significant group differences in the change in latency from baseline (*H*(2) = 1.12, *p =* .57) (Fig. 5I). In short, we observed elevated aggression in terms of bite frequency but not latency in response to alcohol drinking. These differences in aggression are not likely to be attributable to differences in preference for the SuperSac vehicle, as SuperSac vehicle intake (in the last session of access) showed no significant associations with aggression in a subsequent resident-intruder test (the screening session of the following day) in terms of bite frequency (R^2^ = 0.084, *p = .*24) or bite latency (R^2^ = 0.011, *p = .*70) (Fig. 5B).

## Discussion

While acute alcohol intoxication is known to produce heightened aggression, the long-term effects of chronic alcohol intake on both alcohol-involved and alcohol-uninvolved aggression is complex and debated. This is the first preclinical study, to our knowledge, to characterize aggressive phenotypes during protracted abstinence following chronic alcohol drinking. Effective and translational animal models are required to separate the psychopharmacological effects of chronic alcohol intake from the complex non-psychopharmacological factors that confound human research. The present study presents one such model. We applied a highly translational model of voluntary binge drinking to a genetically heterogeneous population, revealing distinct subpopulations which reflect the clinical differences between heavy and moderate alcohol users. We found that repeated cycles of binge-like drinking produce heightened aggression in confrontations that do not involve alcohol. We also found that when animals drink a large amount of alcohol (∼1-1.5 g/kg in a 1 hr session), they are acutely more aggressive regardless of their prior drinking histories.

We adapted a model of repeated binge-like drinking, Drinking in the Dark, to a genetically heterogeneous mouse strain and identified two distinct subpopulations: High Drinkers (average alcohol intake = 1.33 g/kg/h) and Low Drinkers (average alcohol intake = 0.45 g/kg/h). Animals’ alcohol intake stabilized over the first four weeks, after which we began to observe changes in aggressive behavior. We observed a small reduction in alcohol intake on Mondays. This may be attributable to the fact that animals’ cages were changed on Sundays, as cage changes are typically accompanied by a reduction in alcohol intake. Based on BECs collected after the acute alcohol test, we estimated that High Drinkers achieved BECs in the range of ∼70-90mg/dL for each hour of DID. This is a pharmacologically meaningful dose, as the National Institute on Alcohol Abuse and Alcoholism (NIAAA) defines binge drinking as a pattern of alcohol intake that brings blood alcohol concentration (BAC) to 0.08% (80 mg/dL) or higher in about 2 hours in humans (National Institute on Alcohol Abuse and Alcoholism (NIAAA), 2025).We observed similar levels of intake in the acute alcohol test. Furthermore, the levels of alcohol intake that we observed both during DID and during the acute alcohol test were consistent with known aggression-heightening doses (Miczek et al., 1992, 1998), including doses that have been shown to produce heightened aggression following voluntary intake with SuperSac vehicle (Miczek et al., 2022).

Both Low Drinkers and Water Drinkers showed a gradual decline in bite frequency over the course of DID. This was expected, as resident-intruder aggression is a conditioned behavior which requires regular screening in the home cage to maintain stable responding. Furthermore, repeated screening in a neutral arena without intermittent home cage sessions should also contribute to a gradual decline in aggression. This decline, however, did not occur in the High Drinkers. These animals showed the opposite trajectory, with bite frequencies at or above baseline levels over the course of weeks 5 – 8. Latencies were somewhat variable in neutral arena fights. This is likely due to the length of time between test sessions as well as the larger size of the arena. The resident mouse has the opportunity to explore the large arena for 5 s before the intruder is introduced, which can lead to differences in latency based on distance from the intruder mouse.

After one week of abstinence, all animals with an alcohol drinking history initiated a fight within 30 seconds of the first intruder presentation, whereas only about 50% of animals without such a history initiated a fight in that time window. It should be noted that at the time of this fight animals had not been exposed to an intruder in over two weeks. Due to this delay, we expected to see high and variable latencies in the first fight, followed by a reduction in bite latency after repeated daily fights. We saw this expected pattern in the water control group, but not in animals with an alcohol drinking history. Water controls displayed a highly skewed distribution, with some animals initiating a fight immediately, and others not initiating a fight for several minutes. This may be reflective of subpopulation differences. It is possible that the differences in latency that we observed may be indicative of changes in the motivation to fight. Previous research using operant paradigms has shown that acute alcohol intoxication can cause an escalation in the motivation to engage in a fight without affecting fighting performance (Covington et al., 2018). It is possible that similar motivational shifts drive these differences in the latency to initiate a fight while not affecting fighting performance in terms of bite frequency. Future research will investigate the effect of chronic alcohol intake on the motivation to fight using operant paradigms.

We found that a single session of voluntary binge-like alcohol drinking produced heightened aggression regardless of drinking history. Before the acute alcohol test, animals were habituated to consume the sweet SuperSac vehicle solution over three drinking sessions. While there were no significant differences in SuperSac intake during habituation, animals with a history of high binge drinking consumed moderately more SuperSac than the other two groups. It is possible that this reflects alcohol-seeking behavior as the SuperSac solution was presented in the same tubes in which the animals had formerly consumed alcohol during DID. It may also reflect greater sensation or novelty seeking in general, as seen in human binge drinkers (Adan et al., 2017). When alcohol was presented in the SuperSac Vehicle, all groups consumed over 1 g/kg on average within the 1 hr session. The Alcohol-Naïve group achieved moderately lower BECs despite similar levels of intake. This may be related to leakage, differences in metabolism, or differences in their behavioral interactions with the drinking tubes. While we observed escalated aggression across all animals, and no significant differences between groups, we noted that the Alcohol-Naïve animals showed a moderately greater increase in bite frequency compared with the other groups. This may be related to differences in body weight, as they were the heaviest group. It may also reflect a genuinely heightened sensitivity to alcohol’s aggression heightening effects in alcohol naïve animals. We also note that the correlation between BEC and change in bite frequency did not reach significance among Low Drinkers. This may be related to the smaller sample size and wider variability in intake within this group. We did not observe statistically significant changes in bite latency following alcohol intake. However, latencies were already very low at baseline, potentially creating a ceiling effect for latency reduction. We observed, however, a modest reduction in median latency, from 2.5 s to 1 s.

### Relevance to preclinical literature

Our findings also align with and build on existing preclinical literature. Hwa et al. (2015) found that 8 weeks of intermittent access alcohol drinking predicted a greater likelihood of engaging in aggressive behaviors 6-8 hours after the end of alcohol access. Hwa et al. observed differences in the likelihood of engaging in a fight, but not in aggressive performance. In contrast, we were able to detect differences in performance. This may be due to several factors. First, we tested the same animals repeatedly so that changes in aggressive behavior could be observed within subjects over time. Second, we introduced a translational model of binge drinking in which we could identify a subpopulation with a propensity for high drinking. Third, we conducted resident-intruder tests within a neutral arena in order to sensitively detect changes in aggressive behaviors. Finally, the data in the present study goes beyond Hwa et al.’s findings in that we extended the timecourse of abstinence to examine aggression after 24 hrs or 1 week of abstinence and also examined responses to acute alcohol intoxication during abstinence. Notably, Hwa et al. observed escalated aggression after 8 weeks, but not 1 or 4 weeks of alcohol access. Similarly, we observed no changes in aggression after 1 or 4 weeks of DID, but began to observe group differences after 5 weeks which persisted over subsequent weeks. The present study also confirms and builds on Miczek et al.’s (2022) finding that voluntary oral alcohol intake at a dose just over 1g/kg produces heightened aggression. Specifically, Miczek et al. (2022) found that voluntary oral intake of alcohol increases preference for an aggressive encounter over a pro-social, sociosexual alternative. This occurs when animals are given free access to alcohol (consuming 1.2 ± 0.02 g/kg). When the dose was restricted to 1.8 g/kg, mice also increased their responding for an operandum that was reinforced by an aggressive encounter. Notably, animals were given 4 sessions of alcohol self-administration prior to behavioral testing. The authors, therefore, did not examine the effects of voluntary intake in completely alcohol-naïve animals in comparison with experienced alcohol drinkers. Our findings build on this, showing that acute voluntary oral alcohol intake produces heightened aggression in both alcohol-naïve animals and those with chronic drinking histories. Finally, our findings align with previous studies of the effects of DID on social behaviors. Flanigan et al. (2023) found that 3 weeks of DID and 1 week of abstinence produced a decrease in social novelty preference in female but not male B6 mice in a 3 chamber sociability test. Similarly, we observed no changes in social interaction in male CFW mice following 7 weeks of DID and 20 h of abstinence.

### Relevance to clinical literature

Clinical research suggests that individuals undergoing abstinence following chronic alcohol intake show higher levels of anger, hostility, and aggression. However, there are multiple ways to explain this. It may be that (1) prior aggression predicts subsequent heavy drinking, (2) a third variable predicts both high aggression and high drinking, (3) a winner effect arises out of past experiences of fighting while intoxicated, or (4) that a history of chronic alcohol intake directly causes heightened aggression through psychopharmacological mechanisms. Our study provides strong support for hypothesis (4). Our data not only supports hypothesis (4), but also provides some evidence against hypotheses (1), (2), and (3). Firstly, (1) in contrast to human research, which shows consistently that behavioral problems often precede the onset of heavy alcohol use, we did not find strong evidence that prior aggression predicts subsequent binge-like drinking. We observed moderate but nonsignificant trends in this direction. Baseline latencies to bite were moderately predictive of subsequent binge-like drinking. While it is possible that these trends would reach significance in a larger sample, they do not explain the dramatic group differences in aggression that we observed after chronic alcohol intake. Secondly, (2) we also did not find strong evidence that a third variable could explain both alcohol drinking and aggression in our study. We examined body weight, locomotor activity, sociability, and general consummatory behaviors and found no evidence that any of these factors significantly confounded the observed effects. It is possible that there could be hidden variables which we did not examine in the study. However, it should be noted that animals who voluntarily consumed very little or no alcohol (the low drinkers) presented similarly to animals who were assigned to the no-alcohol condition (the H₂O group). Both groups showed a gradual decline in aggression over time with bite frequencies averaging around 10 bites per session in weeks 5-8. In other words, whether animals chose to drink at low levels or were artificially assigned to a no-drinking group, their behavior was categorically distinct from the patterns of aggressive behavior seen in the High Drinkers. This strongly suggests that the observed effects are not due to self-sorting based on a hidden third variable, but rather, are due to the causal effects of chronic alcohol intake. Finally, (3) human research on alcohol-uninvolved aggression following chronic drinking may be confounded by the possibility of a winner effect arising from past experiences of fighting while intoxicated. In other words, it may be that chronic alcohol intake does not cause heightened aggression per-se, but rather, that acute intoxication causes heightened aggression and experiences of aggressive outbursts during intoxication positively reinforce aggressive behavior. This, however, was not a possibility in our study, as animals were not intoxicated during any aggression tests aside from the final test day.

Our data align well with human clinical research showing that different patterns of alcohol intake have different relationships with aggressive behavior. Several studies have shown that heavy, chronic alcohol use, but not moderate drinking, is associated with heightened aggression (Beseler et al., 2012; Kumar Sharma et al., 2012). Furthermore, some evidence suggests that a binge like pattern of intoxication has particularly deleterious consequences, both in general and with respect to aggression (Waterman et al., 2019). Our data suggest that these may well be the case, as we saw distinct patterns of aggression in heavy drinkers compared with moderate drinkers. Additionally, clinical research suggests that aggression, anger, and hostility may present differently in acute compared with protracted abstinence. A study by Fox et al. (2008) found that deficits in emotion regulation persist for only the first few days of abstinence in recovering alcoholics, whereas deficits in impulse control persist for several weeks. Ozsoy & Esel (2008) found that alcoholics show abnormal cortisol non-suppression to dexamethasone during early abstinence, but only individuals who are high in trait aggressivity show this effect during protracted abstinence. Ziherl et al. (2007) found that 3 year abstinent alcoholics show higher levels of indirect aggression and hostility, while differences in physical and verbal aggression disappeared. In line with these findings, we observed distinct aggressive phenotypes during acute compared with protracted abstinence. During acute abstinence, aggressive effects appeared only in High Drinkers and presented as high bite frequencies with no differences in latency. During protracted abstinence, both High and Low drinkers were quicker to initiate their first aggressive encounter and latencies were more consistent in comparison with water controls.

Finally, while our data support the hypothesis that chronic alcohol intake can cause an escalation in alcohol-uninvolved aggression, clinical research suggests that chronic alcohol intake may also be a risk factor for alcohol-involved aggressive incidents (Kingree & Thompson, 2015; Wells et al., 2005; White & Hansell, 1996). A study in Italian adolescents, for example, found that heavy chronic alcohol use is associated with a heightened risk for alcohol-involved aggression, while moderate use is not (Siciliano et al., 2013). This association, however, may be explained in two ways (1) chronic alcohol intake creates a heightened sensitivity to alcohol’s acute aggression-heightening effects or (2) heightened aggression is observed in frequent alcohol users simply because they are more likely to be intoxicated during aggressive encounters.

Our findings support hypothesis (2): that chronic alcohol intake does not significantly alter the acute response to alcohol. We found that a single session of alcohol drinking produced heightened aggression regardless of drinking history. However, it is possible that drinking history may have more subtle effects on the response to acute intoxication. As discussed, Covington et al. (2018) found that a history of repeated alcohol administration produces escalated aggressive responding in response to alcohol in an FI paradigm. Effects were not observed, however, in terms of fighting performance. Given that this study examined fighting performance and not motivation, it remains possible that effects could still be observed in terms of motivation. Future research will need to combine DID with FI paradigms to address this. It is also possible that subtle group differences would be observed in alcohol-heightened aggressive performance in a larger sample using a more consistent method of alcohol delivery. While we may have obtained more consistent aggressive behavioral data had we used a forced method of alcohol administration, such as oral gavage, at a known aggression-heightening dose, our approach of providing an acute alcohol dose via voluntary oral intake has superior face validity (Miczek et al., 2013).

### Potential Mechanisms

Several neural mechanisms may explain the effect of chronic alcohol intake on subsequent alcohol-uninvolved aggression. Prominent models of aggression, such as the Social Information Processing Theory (Crick & Dodge, 1994) and the General Aggression Model (Allen et al., 2018; Anderson et al., 1995) posit that two main factors are responsible for excessive aggression in humans: (1) hostility bias, and (2) poor emotional and impulse control. As stated above, abstinent alcoholics show an exaggerated hostility bias in multiple laboratory paradigms. Abstinent alcoholics also show deficits in emotional regulation and impulse control (Fox et al., 2008; Jakubczyk et al., 2018; Mitchell et al., 2005; Quoilin et al., 2018), and this finding has been replicated in rodent models (Moschak et al., 2013). These two factors may drive reactive aggression through mechanisms involving hyperfunction of the amygdala alongside hypofunction of the mPFC (Flanigan & Russo, 2019). It is possible that chronic, heavy alcohol intake can dysregulate brain function in these regions alongside other regions that are responsible for both threat processing and impulse control, such as the orbitofrontal area, anterior cingulate cortex, as well as subcortical and frontomesal regions (Bartholow & Heinz, 2006; Ochsner & Gross, 2005; Sinha & Li, 2007; Verdejo-García & Bechara, 2009). While the potential neural mechanisms center around long term changes in threat processing and impulse control mechanisms, the relationship between chronic drinking and aggression may also be explained by non-neural mechanisms, such as blunted HPA axis responsiveness (Ozsoy et al., 2006; Shen et al., 2024). Alternatively, aggression can be induced by the frustrative omission of a reward (de Almeida & Miczek, 2002). Therefore, heightened aggression during abstinence may be related to alcohol seeking behavior or anxiety related to the absence of alcohol (Cardoso et al., 2006).

### Future Directions

Future studies might apply this model to examine the neural mechanisms through which chronic alcohol intake alters aggressive behavior. Future studies may examine these mechanisms using high-alcohol drinking mouse strains, such as C57BL/6J mice to produce higher and more consistent BECs, or may use non-voluntary modes of alcohol administration, such as vapor chambers or oral gavage, to better control alcohol dose and eliminate the effects of confounding variables which might contribute to both voluntary alcohol intake and aggression. However, the model that we present here may also prove useful due to its superior face validity. Future studies may examine aggression at time points between 1 – 7 days of abstinence, and may also reintroduce alcohol at a more acute time point during abstinence. We observed notable changes in aggressive behavior after just 20 hours of abstinence. Future studies should reintroduce alcohol at such a time point to examine whether alcohol’s acute aggression heightening effects are more intense at this time.

## Acknowledgements

This research was funded by the National Institutes of Health grants R00AA029726 and R01AA013983 awarded to MMW. We thank Brittany M. Blasetti for her contributions to experimental design and Dr. Randall J. Ellis for his analytical expertise. The authors declare they have no conflicts of interest.

