## Supplement for "Repeated Binge-Like Alcohol Drinking Heightens Aggression in Mice"

#### Fig S1

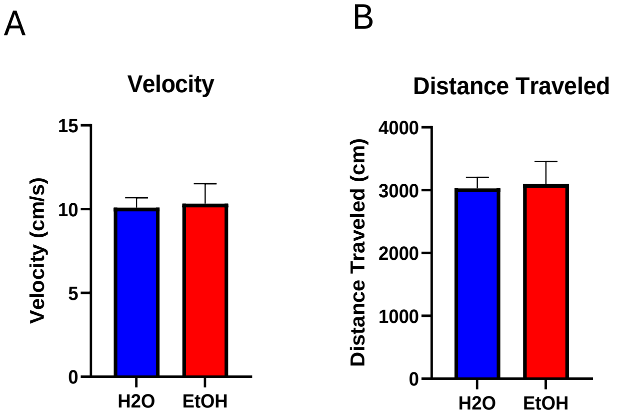


**Fig S1** Prior to neutral arena testing, locomotor activity was measured in an empty plexiglass arena of the same dimensions as the neutral arena. Locomotor activity was assessed over 5 minutes. Prior to beginning Drinking in the Dark, we ensured that there were no differences in either velocity ($p > .05$) (**A**) or distance traveled ($p > .05$) (**B**) between animals assigned to EtOH or $H_2O$ groups. These measures served to confirm that differences in aggression could not be attributed to baseline motor activity in response to a novel environment.

#### Fig S2

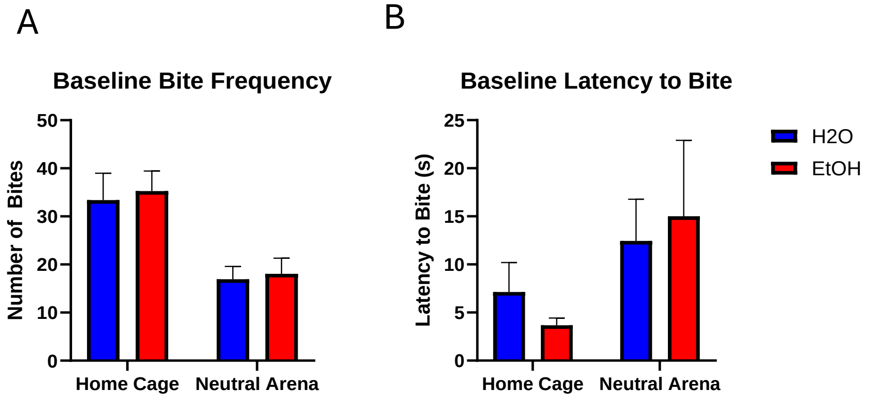


**Fig S2** Baseline aggression was measured prior to the assignment of EtOH or $H_2O$ drinking conditions. Bite frequency (**A**) and latency to bite (s) (**B**) were averaged across the final two screening sessions conducted in both the home cage and neutral arena. Groups were counterbalanced based on baseline bite frequency in the neutral arena and we ensured that there were no statistically significant group differences in any baseline measure of aggression ($p$ > .05).

### Fig S3

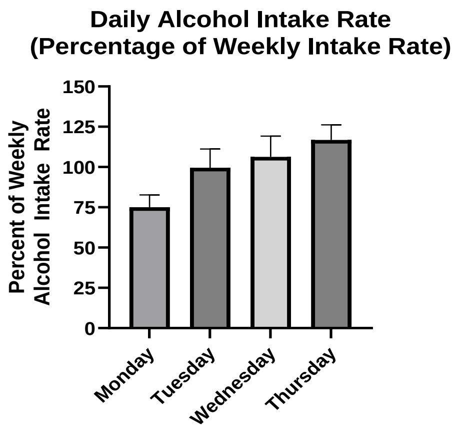


**Fig S3** Average daily alcohol intake increased throughout the week, with the lowest intake observed on Mondays. To account for escalation across repeated DID cycles, intake was normalized as a percentage of each animal's weekly average rate of alcohol intake in g/kg/h. A Friedman test revealed a significant effect of weekday on alcohol normalized alcohol intake rate ($\chi^2$ (3) = 10.30, $p$ = .016).

#### Fig S4

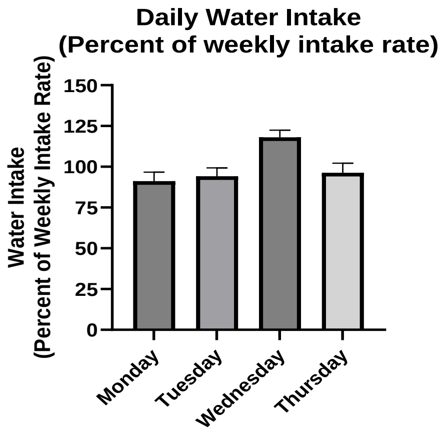


**Fig S4** Average daily water intake peaked on Wednesdays. To account for escalation across repeated DID cycles, intake was normalized as a percentage of each animal's weekly average rate of alcohol intake in ml/h A Friedman test revealed a significant effect of weekday on alcohol normalized alcohol intake rate ($\chi^2$ (3) = 10.30, *p* = .016). A repeated measures ANOVA confirmed that there was a significant effect of weekday on water intake (F (3, 21) = 4.11, *p* = .019), and trend analysis further indicated that water intake followed a cubic pattern (F(1, 7) = 8.22, *p* = .024).

**Fig S5**

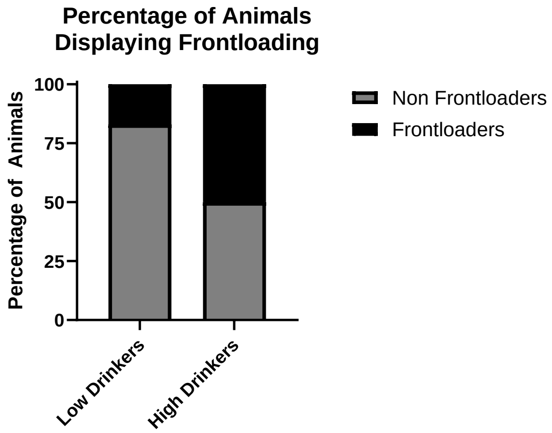


**Fig S5** Frontloading was assessed by comparing the rate of intake (g/kg/min) during the first 20 minutes of each drinking session with the rate of intake in the remainder of the session. Rates were averaged across weeks 5–9 to calculate a “rate ratio” (early vs. late session intake). Animals with a rate ratio greater than 1 were classified as frontloaders. 50% of High Drinkers (3 out of 6) and 17% of Low Drinkers (1 out of 6) were classified as frontloaders.

#### Fig S6

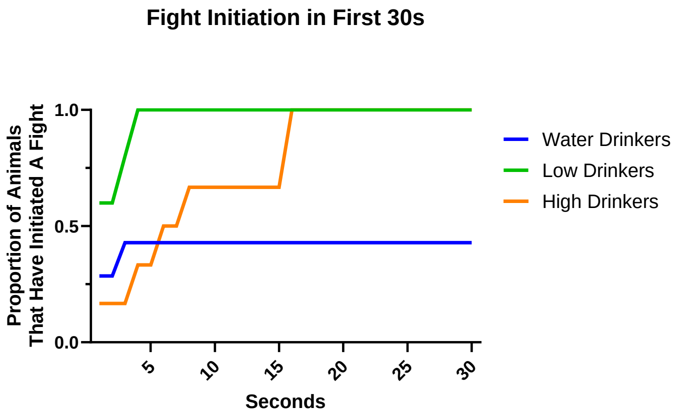


**Fig S6** In the first intruder presentation following one week of abstinence, both Low and High Drinkers rapidly initiated a fight. The proportion of animals that initiated a fight in each second over the first 30 seconds of the test are represented here, showing that all High Drinkers (n = 6) and Low Drinkers (n = 5) initiated a fight within this window, whereas less than 50% of Water Drinkers did so.

**Fig S7**

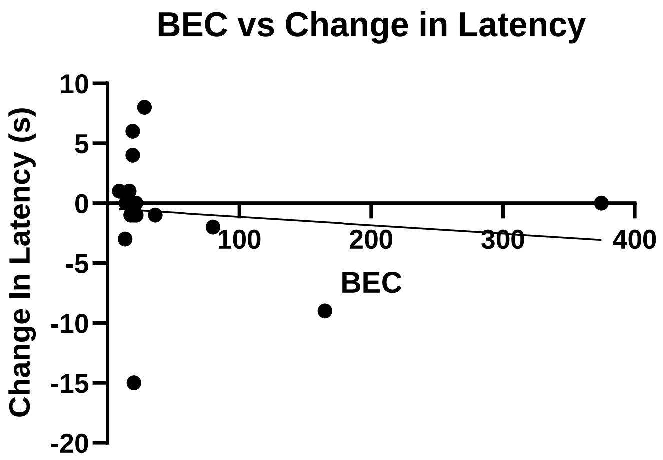


**Fig S7** BECs (mg/dl) were not significantly associated with the change in latency (s) between a baseline resident intruder fight and a fight conducted immediately following a 1 h session of access to 15% EtOH in SuperSac vehicle ($R^2$ = 0.015, p = .65).
